# Roles of TYRO3 Family Receptors in Germ Cell Development During Mouse Testis Formation

**DOI:** 10.1101/2024.03.05.583252

**Authors:** Zhenhua Ming, Stefan Bagheri-Fam, Emily R Frost, Janelle M Ryan, Michele D Binder, Vincent R Harley

## Abstract

**Objective:** To investigate the role of a potential SOX9 target gene, *Tyro3*, along with its family members, *Axl* and *Mertk* (TAM family) in mouse testis development.

**Design:** Experimental laboratory study.

**Setting:** Research institute units.

**Subject(s):** Embryonic day (E)11.5 Swiss mouse gonads for *ex vivo* gonad culture; *Tyro3* knockout mouse embryos.

**Intervention(s):** E11.5 Swiss mouse gonads were cultured in hanging droplets of 30 µL DMEM medium supplemented with 10% FBS and 1% antibiotic-antimycotic. A pair of gonads were treated with 20 μM of BMS-777607 or 30 μM of LDC1267 and an equivalent volume of the vehicle control DMSO.

**Main Outcome Measure(s):** Immunofluorescence to measure morphological changes of *ex vivo* cultured gonads and *in vivo Tyro3* mouse testes; qRT-PCR to measure gene expressions.

**Result(s):** Inhibition of the TAM family in E11.5 *ex vivo* cultured male mouse gonads led to reduced germ cell numbers caused by reduced proliferation and increased apoptosis of the germ cells. *Tyro3* knockout mice exhibited reduced expression levels of the germ cell genes *Ddx4*, *Dazl* and *Pou5f1* and increased expression levels of the Sertoli cell genes *Sox9* and *Amh* at E12.5. However, by E14.5, the expression of *Ddx4*, *Dazl*, *Sox9* and *Amh* had returned to normal levels in *Tyro3* knockout testes. *Tyro3* knockout testes displayed normal morphology and structures during fetal testis development.

**Conclusion(s):** TAM family members have redundant roles in regulating germ cell development during early testis development.

**Attestation Statement:**

- Data regarding any of the subjects in the study has not been previously published unless specified.
- Data will be made available to the editors of the journal for review or query upon request.

**Data Sharing Statement:** **N/A**

**Capsule:** Inhibition of the TAM family led to loss of germ cells in fetal gonads and deletion of *Tyro3* alone disturbed gene expressions of germ cells and Sertoli cells.

## Introduction

In mouse gonadal sex determination, the expression of *Sry* is the driving force behind the formation of testes from the bipotential gonad (1, 2). This initiation leads to the transactivation of *Sox9* in supporting cells, which differentiate into pre-Sertoli cells by embryonic day (E)11.5 (2). Sertoli cells play a central role in testis development, forming testis cords around germ cells and initiating the differentiation of fetal Leydig cells and peritubular myoid cells (3). Conversely, in developing XX gonads, *Sox9* is downregulated, and FOXL2 and WNT4/RSPO1 guide the differentiation of the supporting cells into pre-granulosa cells (4, 5). Between E13.5 and E14.5, XY gonadal germ cells undergo mitotic arrest, while germ cells in the XX gonad enter meiosis triggered by retinoid acid (6, 7).

The balance of SOX9 expression is important for determining male or female gonadal fate. Notably, mutations in human *SOX9* are linked to campomelic dysplasia (CMPD, OMIM# 114290), with around 75% of cases associated with partial or complete XY sex reversal (8–10), underscoring the significance of SOX9 in human testis determination. In mice, conditional homozygous inactivation of *Sox9* in XY gonads leads to complete male-to-female sex reversal. In these *Sox9* knockout gonads, Sertoli cells, testis cords, and Leydig cells are notably absent, and ovarian markers like *Wnt4* and *Foxl2* are upregulated, with the presence of meiotic germ cells (11, 12). These findings establish SOX9 critical role in Sertoli cell differentiation and testis determination in mice, where it plays diverse roles across multiple tissues and organs, regulating downstream genes for various functions (13). Previously, we performed RNA-sequencing on XY *AMH-Cre;Sox9^flox/flox^* knockout gonads to elucidate the role of SOX9 after the sex determination stage (14). Although these mice show normal embryonic testis development, they become infertile by approximately 5 months of age (15), hinting at a potential involvement of SOX9 in germ cell regulation. However, the specific mechanisms remain unknown. We also conducted SOX9 ChIP-sequencing on both bovine and mouse fetal testes, and combined with the RNA-Seq data, we identified a set of potential direct SOX9 target genes in Sertoli cells (14). Among these, *Tyro3* emerged as an intriguing candidate due to its male-specific expression during the early stages of gonad differentiation (16).

TYRO3, together with AXL and MERTK, comprises the TAM family of receptor tyrosine kinases (17, 18). The TAM family is involved in immune regulation, clearance of apoptotic cells, platelet aggregation, and cell proliferation, survival, and migration (19). TAM triple knockout (TAM^−/−^) mice exhibit severe phenotypes, such as male infertility, blindness and a range of autoimmune diseases (20, 21). The degenerative phenotype of infertility observed in TAM triple knockout male mice results from the perturbed clearance of immature and defective spermatids by Sertoli cells, starting at three weeks of age and worsening over time (20, 22). During the early stages of mouse gonad differentiation, TYRO3 appears to be expressed in males only. It is expressed in Sertoli cells of the testis from E11.5 onwards, whereas no TYRO3 signal was observed in ovaries until E17.5 (23). In mouse embryos, *Axl* and *Mertk* are expressed in Sertoli cells and Leydig cells in male gonads, and granulosa cells and stromal cells in female gonads from E11.5 to E13.5 (16). Notably, none of the three receptors are expressed in germ cells.

This study seeks to elucidate the role of TYRO3 and the TAM family in mouse fetal testis development. Our findings reveal that inhibition of the TAM family leads to germ cell loss due to reduced germ cell proliferation and increased germ cell apoptosis in cultured gonads. In addition, examination of *Tyro3* knockout testes reveals decreased expression of germ cell genes and increased expression of Sertoli cell genes at E12.5, despite the absence of morphological abnormalities. These findings emphasize the additive roles for TAM family members in germ cell development.

## Materials and Methods

### Mice

All animal procedures were conducted in accordance with the regulations of the Florey Institute of Neuroscience and Mental Health animal care committee. Experiments using the *Tyro3*^−/−^ were approved by the Animal Ethics Committee at the Florey Institute of Neuroscience and Mental Health and conducted in accordance with the National Health and Medical Research Council (Australia) Guidelines. *Tyro3* knockout (*Tyro3*^-/-^) mice were fully backcrossed onto the C57BL/6J background and maintained under specific-pathogen-free conditions throughout breeding and experimentation, as described previously (24). The original *Tyro3*^-/-^ line was kindly provided by Prof. Greg Lemke from the Salk Institute of Biological Studies, La Jolla, CA (20) and currently maintained by M. Binder at the Florey Institute of Neuroscience and Mental Health. Heterozygous *Tyro3*^+/−^ mice were crossed with each other to obtain homozygous *Tyro3*^−/−^ embryos (denoted as *Tyro3* KO) and *Tyro3*^+/+^ wildtype littermates (denoted as WT) as controls. Embryos were collected at E12.5, E14.5 and E15.5, where the morning of plug identification was designated E0.5. The staging of E12.5 embryos was accurately determined by counting tail somite numbers. Genotyping analysis to identify the *Tyro3* locus and determine the genetic sex using *Smcxy* loci was performed using genomic DNA extracted from tail tissue. Genotyping primer sequences: *Tyro3* mutant (mut): 5’-GCCAGAGGCCACTTGTGTAG-3’; *Tyro3* wildtype (wt): 5’-TCACTGCACCCCTAAGGTTC-3’; *Tyro3* common (com): 5’-CACACACGCAAAATCAGGTC-3’. *Smcxy*-F: 5’-CCGCTGCCAAATTCTTTGG-3’ and R: 5’-TGAAGCTTTTGGCTTTGAG-3’ (25).

### *Ex vivo* gonad culture with TAM family inhibitors

All animal experimentation was approved and carried out according to the guidelines established by the Monash Medical Centre Animal Ethics Committee. The gonad-mesonephros complex was carefully dissected out from E11.5 Swiss mouse embryos and cultured in hanging droplets of 30 µL DMEM medium (Gibco) supplemented with 10% FBS (Bovogen) and 1% antibiotic-antimycotic. A pair of gonads were treated with 20 μM of BMS-777607 (S1561, Selleckchem) (26, 27) or 30 μM of LDC1267 (S7638, Selleckchem) (28) and an equivalent volume of the vehicle control DMSO. Drug concentrations were selected based on previous studies, and their impact on cell viability was assessed in the NT2/D1 testicular cell line using MTS assay, which showed no observable effects (data not shown). The explants were cultured in a humidified cell culture incubator set at 37°C with 5% CO_2_. After 24 hours, gonads were either harvested or the drug treatment was washed off, fresh medium was replaced, and the culture was continued for 24 or 48 hours. Gonads were either fixed in 10% neutral buffered formalin for immunofluorescence staining.

### Immunofluorescence

Immunofluorescence analysis was performed as previously described (29). Primary antibodies used were anti-SOX9 rabbit polyclonal (AB5535, Merck, 1:1000), anti-DDX4 goat polyclonal (AF2030, R&D Systems, 1:500), anti-AMH mouse monoclonal (sc-166752, Santa Cruz, 1:100), anti-Laminin rabbit polyclonal (L9393, Merck, 1:200), anti-phospho-Histone H3 (Ser10) rabbit polyclonal (06570, Millipore, 1:1000), anti-cleaved Caspase-3 rabbit polyclonal (9661, Cell Signaling, 1:1000), anti-GATA4 mouse monoclonal (sc-25310, Santa Cruz, 1:100), and anti-FOXL2 goat polyclonal (ab5096, Abcam, 1:100). The fluorescent-conjugated secondary antibodies (Thermo Fisher Scientific, 1:1000) include donkey anti-rabbit Alexa Fluor 488 (A-21206), donkey anti-mouse Alexa Fluor 594 (A32744), donkey anti-goat Alexa Fluor 594 (A-11058), and donkey anti-goat Alexa Fluor 647 (A32849. Slides were imaged using confocal fluorescence microscopy (FV1200, Olympus Corp). At least two independent gonads per genotype or treatment were examined.

### Quantitative reverse transcription polymerase chain reaction (qRT-PCR)

Total RNA was extracted from fetal gonads (with mesonephros removed) using a RNeasy Mini kit (74104, Qiagen) or a RNeasy Micro kit (74004, Qiagen) according to the manufacturer’s instructions. Complementary DNA (cDNA) synthesis was performed using the QuantiTect Reverse Transcription Kit (205313, Qiagen). qRT-PCR was performed on a QuantStudio™ 6 Flex Real-Time PCR System (Applied Biosystems) using QuantiNova SYBR Green PCR Kit (208054, Qiagen). All primer sequences are shown in **Supplementary Table 1**. Relative gene expression levels were calculated using the delta delta cycle threshold (ΔΔCt) method with *Tbp* as the normalizing control. Statistical significance was determined using the unpaired Student’s *t* test performed in GraphPad Prism 8.

## Results

### TAM family inhibition leads to germ cell loss in fetal male and female gonads

To elucidate the role of the TAM family within the somatic cells during mouse fetal gonad development, E11.5 XY and XX mouse gonads were treated with the TAM family inhibitors BMS-777607 or LDC1267 in *ex vivo* culture. After culture, gonads were sectioned and immunostained with antibodies specific to germ cell marker DDX4, Sertoli cell marker SOX9, somatic cell marker GATA4, or testis cord marker Laminin.

After 72 hours, we observed that the number of DDX4-positive germ cells in XY gonads treated with BMS-777607 was significantly reduced by 10-fold when compared to DMSO-treated XY gonads (**Figure 1A**). To pinpoint the timing of the germ cell defect, we examined the cultured gonads at 24 and 48-hour intervals. After 48 hours, BMS-777607-treated XY gonads already showed 9 times fewer germ cells compared to the control group (**Figure 1A**). However, after 24 hours, the number of germ cells in BMS-777607-treated XY gonads was comparable to control gonads, suggesting that the loss of germ cells occurred between 24 and 48 hours of culture (**Figure 1A**). In female gonads treated with BMS-777607, germ cell depletion was also evident (Supplementary Figure 1A). Compared to the control group, BMS-777607-treated XX gonads showed a moderate loss of germ cells after 48 hours (2.7-fold), whereas after 72 hours the germ cell numbers were reduced by 9-fold, to a similar extent as in the BMS-777607-treated XY gonads. This shows that germ cell loss occurred faster in the male than in the female gonads. Germ cell depletion by TAM family inhibition was confirmed by another TAM family inhibitor LDC1267. Both male and female gonads treated with LDC1267 exhibited a substantial reduction in the number of germ cells (6-fold and 7-fold, respectively), after 72 hours of culture, as evidenced by DDX4 immunostaining (**Figure 1B** and Supplementary Figure 1B). Taken together, these results highlight the requirement of the TAM family for germ cell development in both male and female gonads.

**Figure 1.**
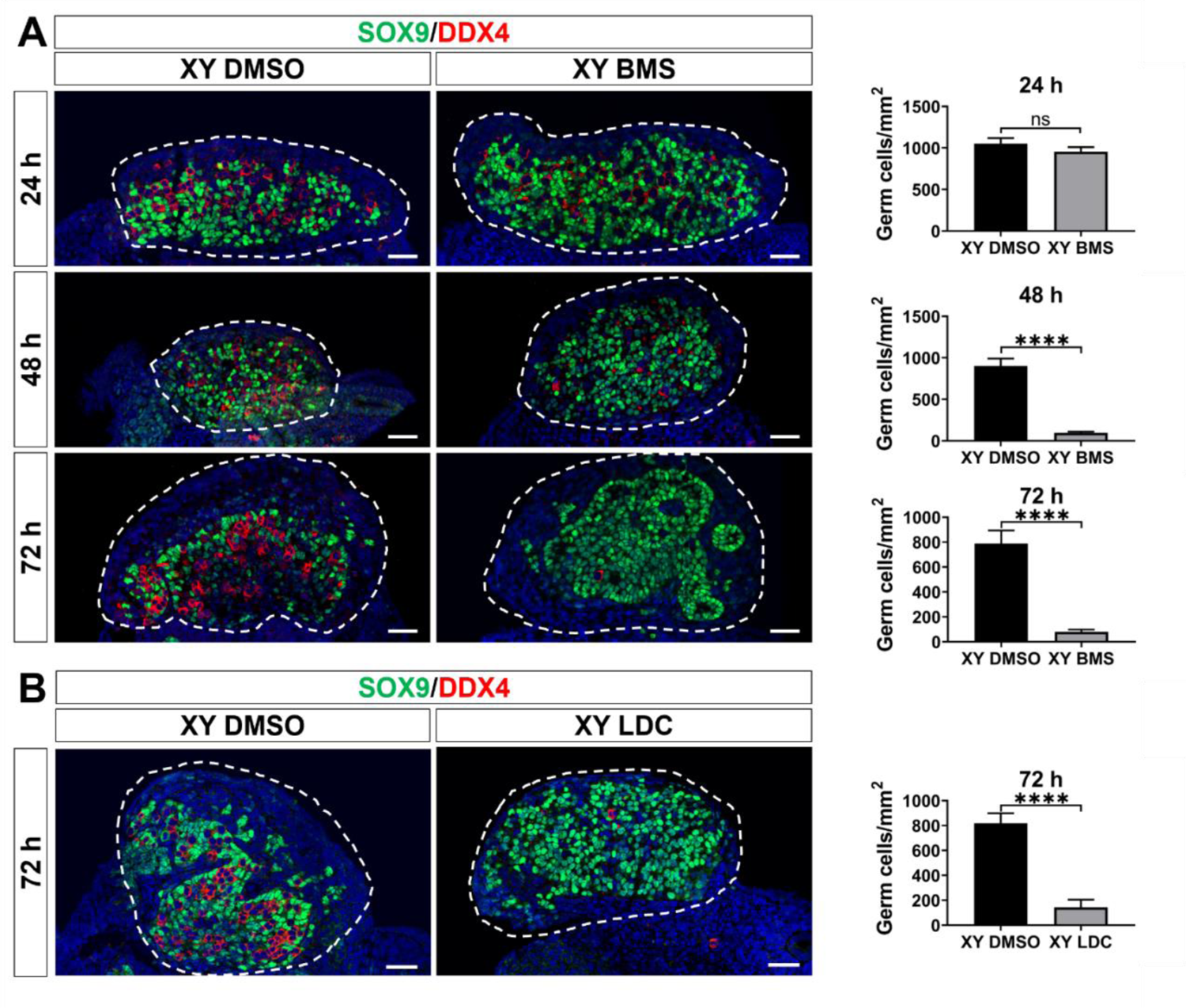
TAM family inhibition induces germ cell loss in XY gonads. (**A**) Immunofluorescence staining of XY gonads treated with either the vehicle control DMSO or the TAM family inhibitor BMS-777607 (BMS) for 24 hours and cultured *ex vivo* for 24, 48, and 72 hours. (**B**) Immunofluorescence staining of XY gonads treated with the TAM family inhibitor LDC1267 (LDC) for 24 hours and cultured *ex vivo* for 72 hours. The sections are stained for the germ cell marker DDX4 (red) and the Sertoli cell marker SOX9 (green). Nuclei are visualized with the nuclear marker DAPI (blue). Scale bar = 50 μm. Dashed lines outline the gonads. Germ cell quantification is determined based on the number of germ cells relative to the area (mm²). At 24 hours, n = 5 for XY DMSO, n = 2 for XY BMS. At 48 hours, n = 3 for XY DMSO, n = 4 for XY BMS. At 72 hours, n = 3 for XY DMSO, n = 4 for XY BMS, and n = 4 for XY DMSO, n = 3 for XY LDC. Data is represented as Mean ± SEM. Unpaired Student’s *t* test. \*\*\*\**P* < 0.0001; ns, not significant.

We next analysed the impact of TAM family inhibition on Sertoli cell development. After 72 hours of *ex vivo* culture, the number of SOX9-positive Sertoli cells in BMS-777607-treated XY gonads was slightly increased by 1.5-fold when compared to control gonads (Supplementary Figure 2A). Sertoli cells exhibited a more densely packed arrangement compared to the control gonads where Sertoli cells typically formed a circular configuration enveloping the germ cells (**Figure 1A**). This may be caused by the reduced number of germ cells in the BMS-777607-treated XY gonads.

Integrity of testis cord was analysed by staining of laminin, a major component of the basal lamina secreted by Sertoli cells and peritubular cells. Like the control XY gonads, BMS-777607-treated XY gonads exhibited the formation of testicular cords, clearly delineated by a laminin-positive basal lamina (Supplementary Figure 3). Similar to the BMS-777607-treated XY gonads, XY gonads treated with LDC1267 showed a more densely packed arrangement of Sertoli cells compared to the control gonads, due to the severe reduction in the number of germ cells (**Figure 1B**). However, in contrast to BMS-777607, TAM family inhibition with LDC1267 did not lead to a significant increase in the number of Sertoli cells (Supplementary Figure 2B). Taken together, these results suggest that Sertoli cell differentiation and testis cord formation is not affected by the loss of the TAM family, despite the loss of germ cells.

### Germ cell number reduction due to reduced proliferation and increased apoptosis

To understand the underlying causes contributing to the near complete loss of germ cells in XY gonads following TAM inhibitor treatment, we investigated cell proliferation and cell apoptosis via immunofluorescence in cultured gonads using phospho-histone H3 (PH3) and cleaved caspase-3 (CC3), respectively. To detect germ cells and somatic cells we also included antibodies for DDX4 or GATA4, respectively. Analyses were done after 24 hours of culture when inhibitor-treated gonads still contained large numbers of germ cells. In control XY gonads, 26.7% of germ cells were proliferating, whereas the percentage of proliferating germ cells in XY gonads treated with BMS-777607 was only 5% (5.3-fold reduction) (**Figure 2A**). Moreover, germ cells in XY gonads treated with BMS-777607 showed a significant 8.3-fold increase in apoptosis when compared to control gonads (13.9% versus 1.7%) (**Figure 2B**). Taken together, these results indicate that the reduction in the number of germ cells in the presence of TAM family inhibitors is attributed to both reduced germ cell proliferation and increased apoptosis.

**Figure 2.**
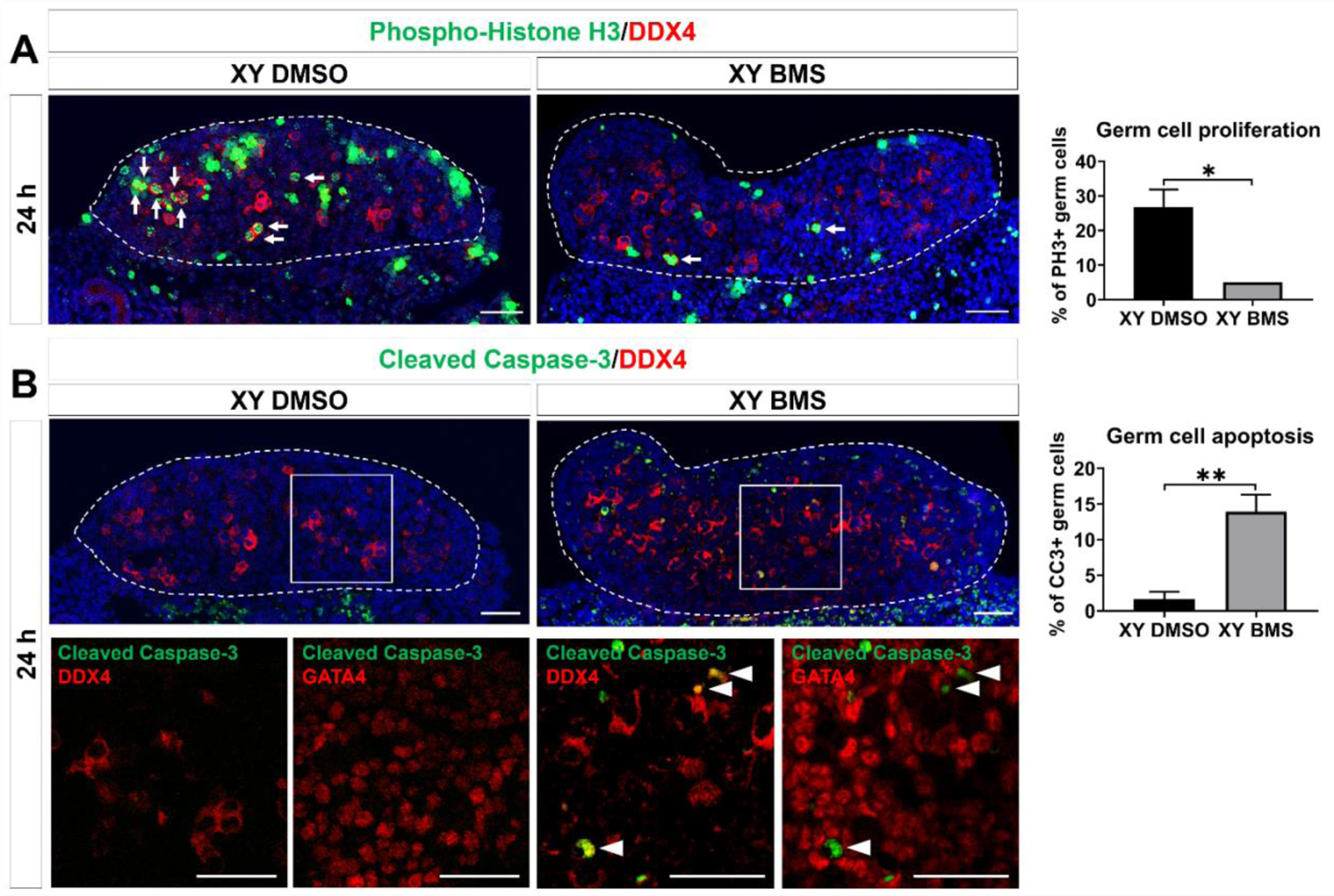
TAM family inhibition leads to decreased germ cell proliferation and increased germ cell apoptosis. (**A**) Proliferation analysis at 24 hours in XY gonads treated with DMSO or the TAM family inhibitor BMS-777607 (BMS) by immunofluorescence staining for the cell proliferation marker phospho-histone H3 (PH3) (green) and the germ cell marker DDX4 (red). White arrows indicate proliferating germ cells. Quantification is performed for PH3+ germ cells relative to the total germ cell population. n = 3 for XY DMSO and n = 2 for XY BMS. (**B**) Apoptosis analysis at 24 hours in XY gonads treated with DMSO or BMS by immunofluorescence staining for the cell apoptosis marker cleaved Caspase-3 (CC3) (green), the germ cell marker DDX4 (red), and the somatic cell marker GATA4 (red). Arrowheads indicate apoptotic germ cells. Quantification is performed for CC3+ germ cells relative to the total germ cell population. n = 4 for XY DMSO and n = 2 for XY BMS. Nuclei are visualized with the nuclear marker DAPI (blue). Scale bar = 50 μm. Dashed lines outline gonads. Data is represented as Mean ± SEM. Unpaired Student’s *t* test. \**P* < 0.05, \*\**P* < 0.01.

### *Tyro3* knockout mice exhibit altered expression of germ cell genes and Sertoli cell genes

Given that *Tyro3* displays male-specific expression and is identified as a potential SOX9 target gene during gonadal development, we next examined *Tyro3* in male gonads in more detail. Gonads were collected from wildtype (WT) and *Tyro3* single knockout (KO) mice at E12.5, shortly after the initiation of sex determination, and at E14.5, when male germ cells enter mitotic arrest while female germ cells initiate meiosis. The expression levels of germ cell-related genes, testis and ovary-specific markers, as well as putative TYRO3 target genes were measured by qRT-PCR.

At E12.5, the expression levels of the germ cell genes *Ddx4*, *Dazl*, and *Pou5f1* were significantly reduced in XY *Tyro3* KO testes compared to XY WT testes (**Figure 3A**). In contrast, the expression levels of germ cell pluripotency markers—*Sox2*, *Nanog*, and *Nanos3*—remained unchanged between XY WT and XY *Tyro3* KO gonads (Supplementary Figure 4A). Next, we analysed gene expression of the Sertoli cell marker *Sox9*, and its direct target genes, including *Amh*, *Dhh*, *Sox10* and *Ptgds* in Sertoli cells. Notably, *Sox9* expression was increased in XY *Tyro3* KO gonads relative to XY wildtype gonads, with *Amh* also showing upregulation in *Tyro3* KO testes (**Figure 3B**). In contrast, the other three SOX9 target genes, *Sox10*, *Dhh*, and *Ptgds* showed no significant differences in XY *Tyro3* KO gonads compared to XY controls, although *Sox10* and *Dhh* showed a trend toward upregulation (**Figure 3B** and Supplementary Figure 4B). The pro-ovarian marker *Foxl2* remained unaffected upon *Tyro3* ablation in XY gonads, with expected higher expression in XX wildtype ovaries (Supplementary Figure 4C).

**Figure 3.**
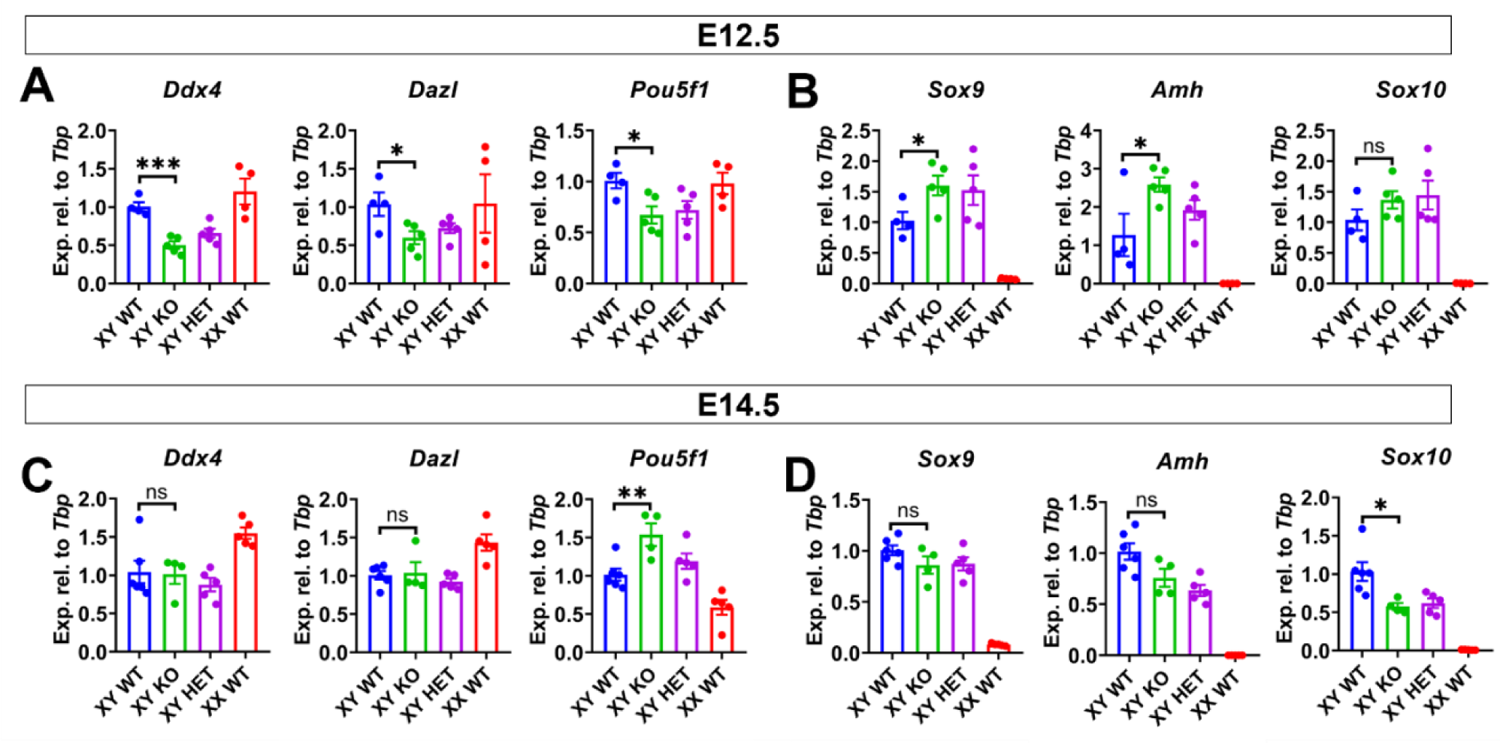
Gene expression profiles of germ cells and Sertoli cells in *Tyro3* knockout testes at E12.5 and E14.5. (**A**, **B**) At E12.5, qRT-PCR analysis of germ cell markers and Sertoli cell markers in XY wildtype (XY WT, blue, n = 4), XY *Tyro3* knockout (XY KO, green, n = 5), XY heterozygous (XY HET, purple, n = 5) and XX wildtype (XX WT, red, n = 4). (**C**, **D**) At E14.5, qRT-PCR analysis of germ cell markers and Sertoli cell markers in XY wildtype (XY WT, blue, n = 6), XY *Tyro3* knockout (XY KO, green, n = 4), XY heterozygous (XY HET, purple, n = 5) and XX wildtype (XX WT, red, n = 5). Gene expression levels are normalized to *Tbp* and presented as relative expression. Data is represented as Mean ± SEM. Unpaired Student’s *t* test. \**P* < 0.05, \*\**P* < 0.01, \*\*\**P* < 0.001; ns, not significant.

At E14.5, the expression levels of *Ddx4* and *Dazl* showed no significant differences between XY WT and KO testes (**Figure 3C**), indicating that their expression recovered in XY *Tyro3* KO testes by E14.5. *Pou5f1* expression was even increased in XY KO testes compared to XY WT testes at E14.5 (**Figure 3C**), also recovering from the downregulation observed in XY KO testes at E12.5. Like at E12.5, the expression levels of *Nanos3*, *Nanog*, and *Sox2* showed no significant differences between XY WT and XY *Tyro3* KO gonads at E14.5 (Supplementary Figure 4F). Unlike E12.5 *Tyro3* KO testes which showed upregulation of *Sox9* and its direct target *Amh*, E14.5 XY *Tyro3* KO testes showed unchanged expression levels of *Sox9* and *Amh* (**Figure 3D**). Notably, *Sox10* expression was significantly reduced in XY *Tyro3* KO testes (**Figure 3D**).

To investigate potential compensatory effects from the other two TAM family receptors, *Axl* and *Mertk*, in *Tyro3*-deficient mouse testes, we analysed *Axl* and *Mertk* in male gonads. *Mertk* expression was reduced in *Tyro3* KO testes at E14.5, while it remained unchanged at E12.5 (Supplementary Figure 4D, I). *Axl* gene expression remained unchanged in XY *Tyro3*-deficient testes compared to control testes at E12.5 and E14.5 (Supplementary Figure 4D, I). Lastly, we questioned whether *Tyro3* ablation in mouse testes could perturb the TYRO3 signaling pathway. Specifically, we aimed to examine any alterations in the expression of key genes associated with the TYRO3 signaling cascade, including *Ccnd1*, *Map2k1*, *Bcl2l11*, *Dusp6* and *Etv5* (30, 31). Among the five candidate TYRO3 target genes, only *Dusp6* was upregulated in *Tyro3* knockout testes compared to wildtype testes at E12.5 (Supplementary Figure 4E). In contrast, *Bcl2l11* was downregulated in E14.5 *Tyro3* KO testes (Supplementary Figure 4J).

In summary, these data suggest that loss of *Tyro3* in Sertoli cells leads to diminished expression of germ cell markers alongside increased expression of Sertoli cell markers at E12.5, whereas these changes restored as gonad development progressed by E14.5.

### Lack of gross morphological changes in *Tyro3* knockout testes

To determine whether *Tyro3* knockout testes display gross morphological changes during development, we conducted immunofluorescence on mouse testes at E12.5, E14.5, and E15.5. Double immunofluorescence for the Sertoli cell marker SOX9 and the germ cell marker DDX4 revealed that Sertoli cells surrounded germ cells in a tubular arrangement, and no differences in testicular structure or size were observed between WT and KO testes (**Figure 4A**). Furthermore, the basal lamina of testicular cords within the KO testes exhibited Laminin-positive demarcation like WT testes, segregating the testicular cords from the adjacent interstitial tissue (**Figure 4B**). The expression of the Sertoli cell differentiation marker AMH in mutant testes was similar to that of controls between E12.5 and E15.5 (Supplementary Figure 5A). This suggests that *Tyro3* knockout mice did not show a delay in testis development. Immunostaining of the ovarian marker FOXL2 revealed no ectopic feminization within the *Tyro3* knockout testes (Supplementary Figure 5B).

**Figure 4.**
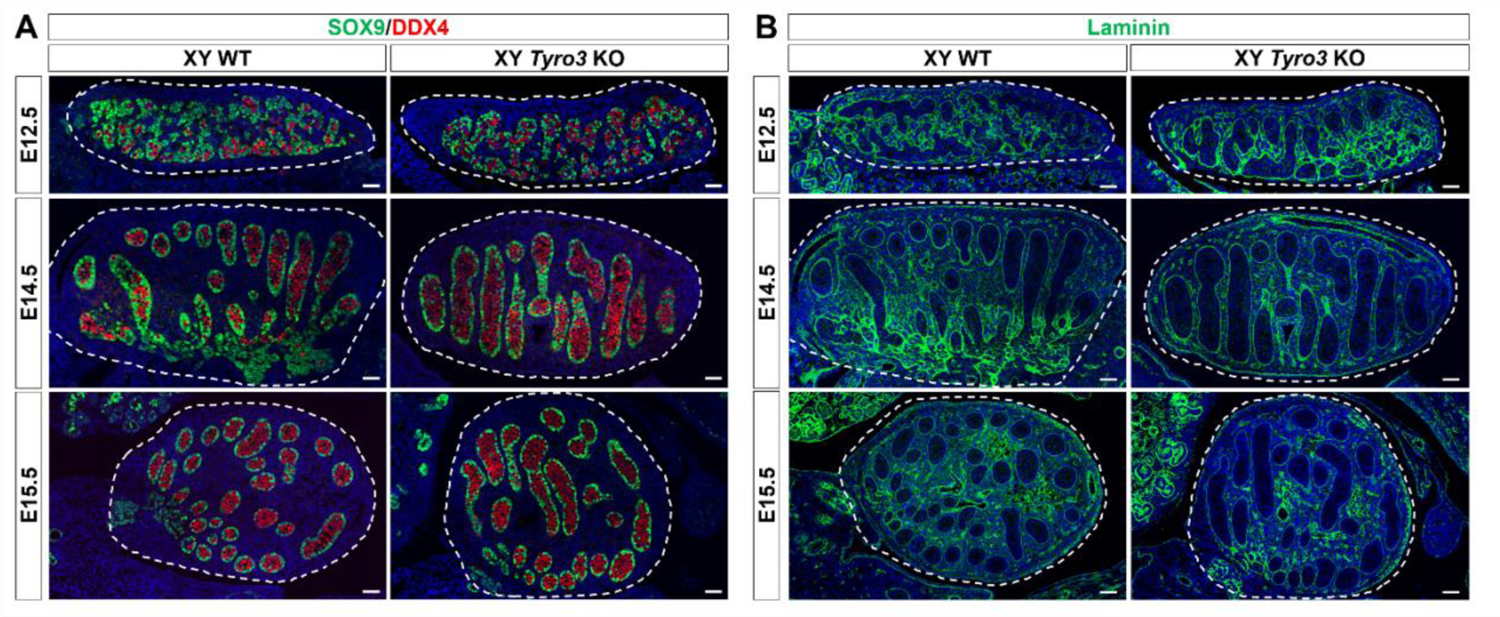
Unaltered fetal testis development in *Tyo3* knockout mice. (**A**) Immunofluorescence staining for the Sertoli cell marker SOX9 (green) and the germ cell marker DDX4 (red), and (**B**) the basal lamina marker Laminin (green) at E12.5, E14.5 and E15.5. Nuclei are visualized with the nuclear marker DAPI (blue). Scale bar = 50 μm. Dashed lines outline gonads.

To analyze cell proliferation and apoptosis within germ cells and Sertoli cells, we examined the expression of the mitotic cell marker PH3 and the cell apoptosis marker cleaved Caspase-3 along with the germ cell marker DDX4 and the somatic cell marker GATA4 at E12.5, E14.5, and E15.5. At E14.5, *Tyro3* knockout testes showed a 12% increase in germ cell proliferation and a 10% increase in Sertoli cell proliferation relative to XY controls (Supplementary Figure 6). However, germ cell and Sertoli cell proliferation levels were similar between *Tyro3* wildtype and mutant testes at E12.5 and E15.5. Furthermore, immunostaining for Caspase-3 showed very little cell apoptosis across the testes in both wildtype and knockout mice (Supplementary Figure 7).

## Discussion

SOX9 is a pivotal transcription factor in testis development, but its downstream target genes and their respective roles have remained largely unexplored. SOX9 orchestrates the development of various organs and tissues through its regulation of target genes (13). In a previous study, we identified potential SOX9 downstream targets during testis development (14). Here, we explore the hypothesis that *Tyro3*, one of these potential SOX9 target genes, and its TAM family members, *Axl* and *Mertk*, mediate SOX9 functions in embryonic testis development. Our findings indicate that inhibiting these TAM family receptors results in a reduction of the germ cell population in both male and female gonads. This reduction is associated with decreased proliferation and increased apoptosis in germ cells, emphasizing the essential role of the TAM family in somatic cells for germ cell development in both testes and ovaries. The observed faster loss of germ cells in male when compared to female gonads following TAM family inhibition aligns with a previous *in vivo* study of TAM triple knockout mice, reporting primary infertility in all male knockout mice while several female knockout mice retained fertility (20). This discrepancy may be attributed to the male-specific and SOX9-driven expression of *Tyro3* in Sertoli cells from E11.5 to E13.5 (14, 16). Further investigation into *Tyro3* single knockout male mice revealed an upregulation of Sertoli cell genes and downregulation of germ cell genes at E12.5, suggesting that TYRO3 plays a role in male germ cell development and may contribute, at least in part, to male infertility in *Sox9* KO mice. However, these gene expression changes resolved by E14.5, and in contrast to TAM family inhibition in *ex vivo* gonad cultures, no abnormalities were detected in fetal testis development in *Tyro3* knockout mice, suggesting redundant functions within the TAM family. This redundancy aligns with observations in TAM family triple knockout males being infertile, whereas the single knockouts show normal fertility parameters (20, 32). Analyzing fertility in these single knockout mice at later stages could reveal potential roles in spermatogenesis, as initially fertile males can develop severe spermatogenesis defects during adulthood (15). For example, *AMH-Cre;Sox9^flox/flox^* conditional knockout male mice become infertile only after 5 months (15).

Our results illuminate the cell non-autonomous influence of TYRO3, AXL, and MERTK, expressed in somatic Sertoli and Leydig cells, on germ cells. This indirect effect appears to involve signals that enhance germ cell proliferation, underscoring the intricate interplay of signals between somatic cells and germ cells in testis development. Sertoli cells function as the organizing center of testis development, releasing various signals that impact other cell types, including germ cells and Leydig cells in the testis (3). For example, FGF9, a SOX9 target expressed in Sertoli cells, promotes germ cell survival in the fetal testis (33). Another SOX9 target gene, *Cyp26b1* in Sertoli cells, antagonizes retinoic acid, preventing male germ cells from entering meiosis (6). In a previous study, we observed the influence of somatic cells on germ cell development in fetal ovaries, where *Fgfr2* deletion in somatic cells led to germ cell loss in ovaries (34). Our findings underscore the significant non-autonomous influence of TAM family members on germ cell development.

Germ cells genes *Ddx4*, *Dazl*, and *Pou5f1* were significantly reduced in *Tyro3* KO testes at E12.5, suggesting a role for TYRO3 in early germ cell development. However, the expression of *Ddx4* and *Dazl* was restored in *Tyro3* KO testes by E14.5, indicating that TYRO3 does not affect their expression after they undergo mitotic arrest. By E14.5, in contrast, *Pou5f1* expression increased in *Tyro3* KO testes, accompanied by increased germ cell proliferation, suggesting compensatory mechanisms for maintaining normal male fertility. Possible mechanisms for the recovery of germ cell development in *Tyro3* KO testes include the potential compensatory roles of *Axl* and *Mertk*, as TAM triple knockout mice were reportedly infertile, whereas single knockout of TAM family members did not impact mouse testis development (20, 32). Additionally, germ cells may possess inherent recovery functions, allowing them to overcome subtle changes in gene expression without affecting overall germ cell development (35). Moreover, the robust reproductive capacity of mice may contribute to germ cell recovery, even in the presence of significant gene expression changes.

*Tyro3* was initially identified as a SOX9 target due to its downregulation in *Sox9* knockout testes at E13.5, supported by ChIP-seq data indicating SOX9 binding to the *Tyro3* promoter (14). When we examined *Sox9* and other SOX9 targets in Sertoli cells of *Tyro3* knockout testes at E12.5, we observed an upregulation of *Sox9* and *Amh*, with *Sox10* and *Dhh* showing a tendency towards upregulation. This might be attributed to a disrupted SOX9-TYRO3 negative feedback loop, resulting in higher *Sox9* levels compensating for *Tyro3* loss and subsequently upregulating SOX9 target genes *Amh*, *Dhh*, and *Sox10*. A transient increase in Sertoli cell proliferation was observed in E14.5 *Tyro3* knockout testes, potentially linked to the elevated SOX9 levels present at E12.5. Notably, *Sox10* was downregulated in *Tyro3* knockout testes at E14.5. SOX10, a target of SOX9, may also have a direct relationship with *Tyro3* in the testis. A previous study demonstrated that TYRO3 positively regulates SOX10 nuclear localization in melanoma cells (36). It is plausible that in Sertoli cells, TYRO3 regulates SOX10 nuclear localization, a hypothesis that warrants further investigation.

TYRO3 downstream genes are implicated in cell cycle progression and anti-apoptosis via the MAPK/ERK and PI3K/Akt signaling pathways (19, 30). In cancer cells, *Tyro3* depletion is linked to downregulation of MAPK pathway components (*MAP2K1*, *MAP2K3* and *MAPK14*), cell cycle progression genes (*CCND1* and *CCND2*), and upregulation of pro-apoptotic genes (*APAF1*, *CASP9*, and *BCL2L11*). *Dusp6* and *Etv5* are components of the fibroblast growth factor receptor (FGFR) tyrosine kinase signaling pathway (31). Activated by FGFR signaling, DUSP6 serves as an *in vivo* negative feedback regulator of the FGFR and MAPK signaling pathways (37). We analysed the expressions of *Ccnd1*, *Map2k1*, *Bcl2l11*, *Dusp6* and *Etv5* in *Tyro3* knockout testes. Altered expression of *Dusp6* and *Bcl2l11* was observed in *Tyro3* knockout testes compared to wildtype, but interpretation is complex due to gene expression in multiple gonadal cell types (16), potentially masking specific effects in supporting cells. As previously mentioned, E14.5 *Tyro3* knockout testes displayed increased expression of *Pou5f1*, potentially contributing to increased germ cell proliferation and explaining the reduced expression of pro-apoptosis marker *Bcl2l11* in *Tyro3* knockout testes.

We observed a downregulation of *Mertk* in *Tyro3* knockout testes at E14.5. In the retinal pigment epithelium (RPE), *Tyro3* expression was reduced in *Mertk* mutant RPE cells, while *Mertk* maintained wildtype expression levels in *Tyro3* mutant RPE cells (38). Notably, *Tyro3* and *Mertk* both reside on chromosome 2 in mice. Previous studies demonstrated alterations in gene expression on chromosome 2 in *Mertk*^-/-^ RPE, including changes in *Tyro3* expression (39, 40). Hence, these findings suggest that within the context of mouse testes, *Tyro3* may indeed influence the expression of *Mertk*.

While our studies provide valuable insights into TAM family roles in germ cell and Sertoli cell development, certain limitations exist. Analyzing whole gonads poses challenges in distinguishing specific cell types affected by TAM family inhibition or *Tyro3* depletion. Future studies utilizing fluorescent transgenic mouse models for sorting and analyzing distinct cell types could enhance understanding of TAM family roles within the testis. The absence of a significant testis phenotype in *Tyro3* single gene knockout mice suggests that *Tyro3* plays no critical role in sex determination. However, it is possible that gonadal development was compromised to some extent, but only mildly and/or transiently. Crossing *Tyro3* knockout mice with partially compromised strains, such as *Sox9* heterozygous mutant mice, may reveal a more robust phenotype (11, 41).

## Conclusions

In conclusion, inhibition of the TAM family results in germ cell loss due to reduced germ cell proliferation and increased germ cell apoptosis, indicating the requirement of the TAM family for germ cell development in fetal gonads. TYRO3 alone affects the expression of germ cell genes and Sertoli cell genes in fetal testes, although no morphological disruptions. Further research is necessary to unravel the intricate gene regulatory network downstream of SRY and SOX9 as well as functions of individual TAM family members.

## Acknowledgements

The authors acknowledge the Monash Medical Centre Animal Facility, the Medical Genomics Facility at the Monash Health Translation Precinct (MHTP), the MHTP Monash Micro Imaging, and the MHTP Monash Histology Platform.

## Author’s contributions

ZM: Writing–original draft, Writing – review and editing, Conceptualization, Investigation, Methodology, Formal Analysis, Project administration, Visualization. SB-F: Writing – review and editing, Conceptualization, Supervision, Data curation. ERF: Writing – review and editing, Supervision. JMR: Writing – review and editing, Methodology. MB: Writing – review, Methodology. VRH: Writing – review and editing, Funding acquisition, Resources, Conceptualization, Project administration, Supervision.

## Supplementary data

**Table S1.**
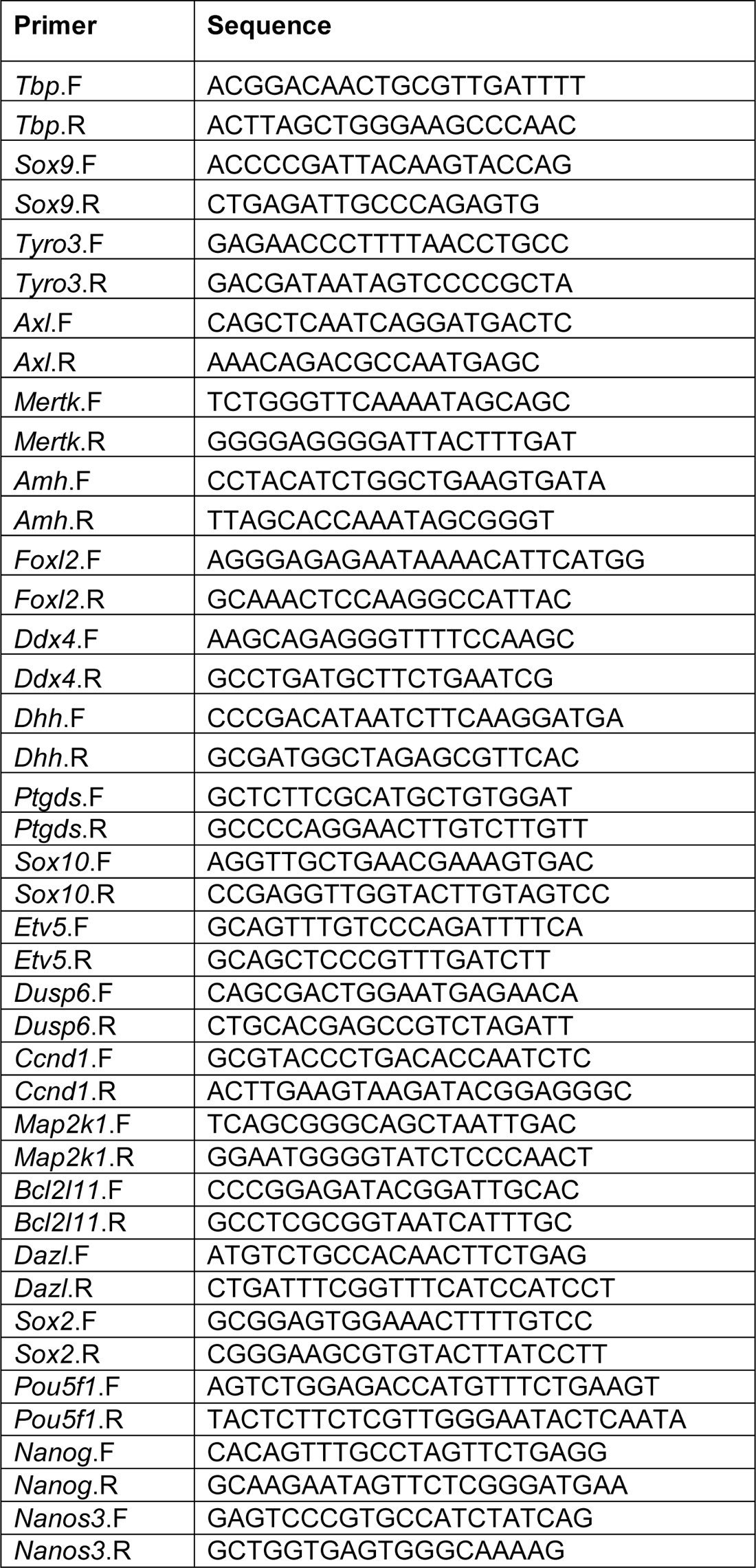
Mouse primer sequences used in qRT-PCR.

**Supplementary Figure 1.**
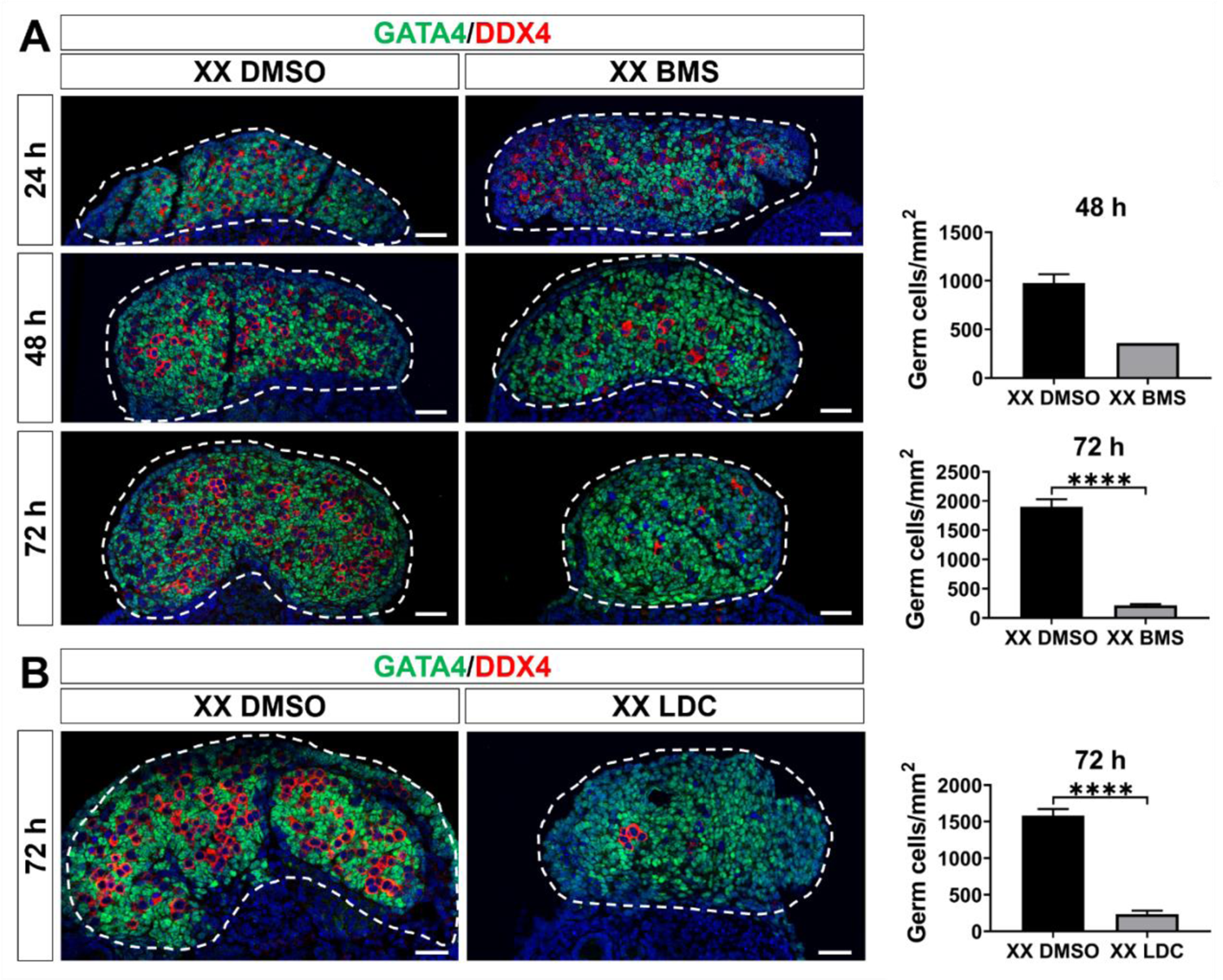
TAM family inhibition induces germ cell loss in XY gonads. (**A**) Immunofluorescence staining of XX gonads treated with either the vehicle control DMSO or the TAM family inhibitor BMS-777607 (BMS) for 24 hours and cultured *ex vivo* for 24, 48, and 72 hours. (**B**) Immunofluorescence staining of XX gonads treated with the TAM family inhibitor LDC1267 (LDC) for 24 hours and cultured *ex vivo* for 72 hours. The sections are stained for the germ cell marker DDX4 (red) and the somatic cell marker GATA4 (green). Nuclei are visualized with the nuclear marker DAPI (blue). Scale bar = 50 μm. Dashed lines outline the gonads. Germ cell quantification is determined based on the number of germ cells relative to the area (mm²). At 48 hours, n = 2 for XX DMSO, n = 1 for XX BMS. At 72 hours, n = 4 for XX DMSO, n = 3 for XX BMS, and n = 9 for XX DMSO, n = 10 for XX LDC. Data is represented as Mean ± SEM. Unpaired Student’s *t* test. \*\*\*\**P* < 0.0001; ns, not significant.

**Supplementary Figure 2.**
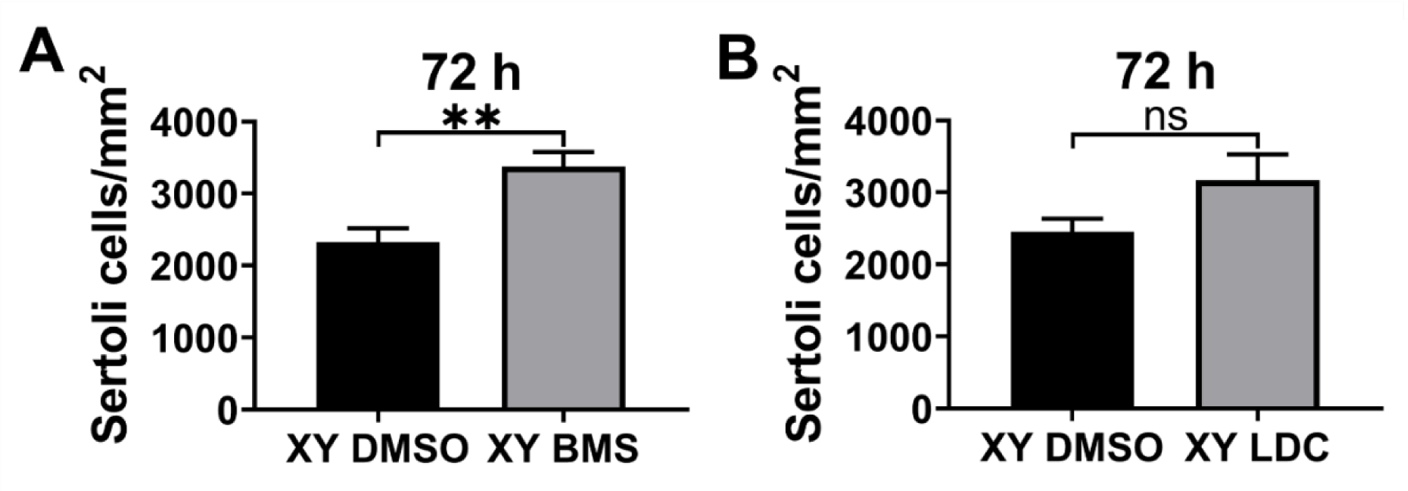
Impact of TAM family inhibition on Sertoli cells in XY gonads at 72 Hours. (**A**) Quantification of Sertoli cells in XY gonads treated with the vehicle control DMSO or the TAM family inhibitor BMS-777607 (BMS) or (**B**) LDC1267 (LDC) for 24 hours and cultured *ex vivo* for 72 hours. Sertoli cell quantification is determined based on the number of Sertoli cells relative to the area (mm²). n = 3 for XY DMSO, n = 4 for XY BMS. n = 6 for XY DMSO, n = 5 for XY LDC. Data is represented as Mean ± SEM. Unpaired Student’s *t* test. \*\**P* < 0.01.

**Supplementary Figure 3.**
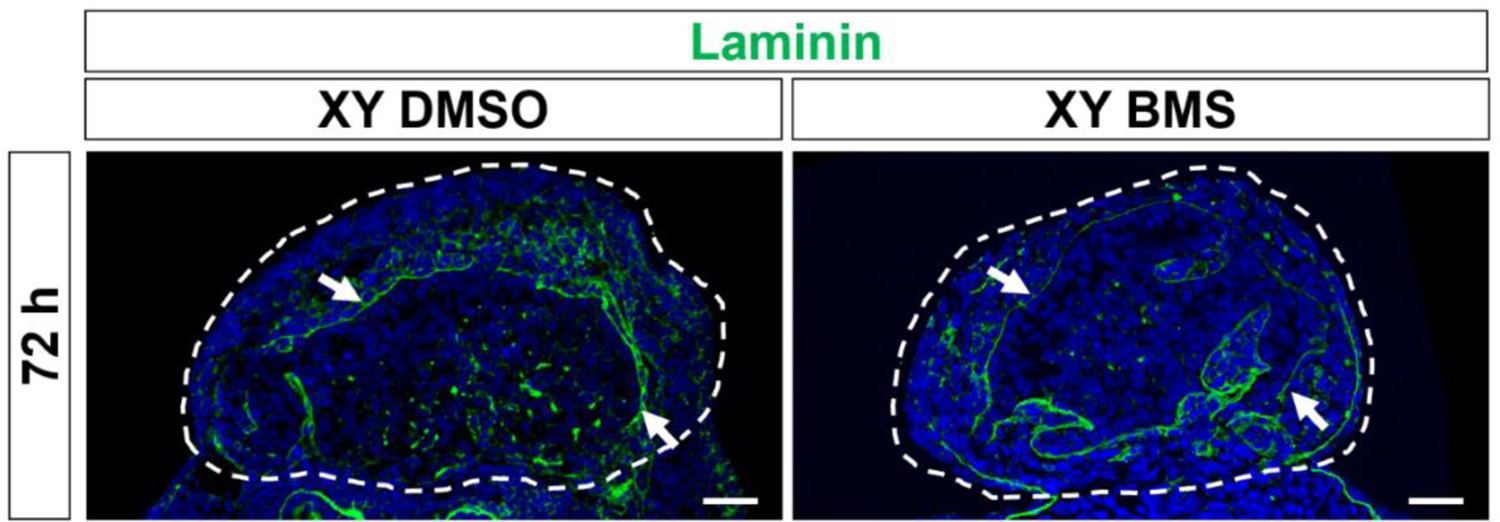
Effect of TAM family inhibition on testis cord formation. Immunofluorescence staining for the basal lamina marker Laminin (green) in control (XY DMSO) and BMS-777607-treated (XY BMS) gonads cultured *ex vivo* for 72 hours. Nuclei are visualized with the nuclear marker DAPI (blue). Scale bar = 50 μm. Dashed lines outline gonads. White arrows indicate intact testis cords.

**Supplementary Figure 4.**
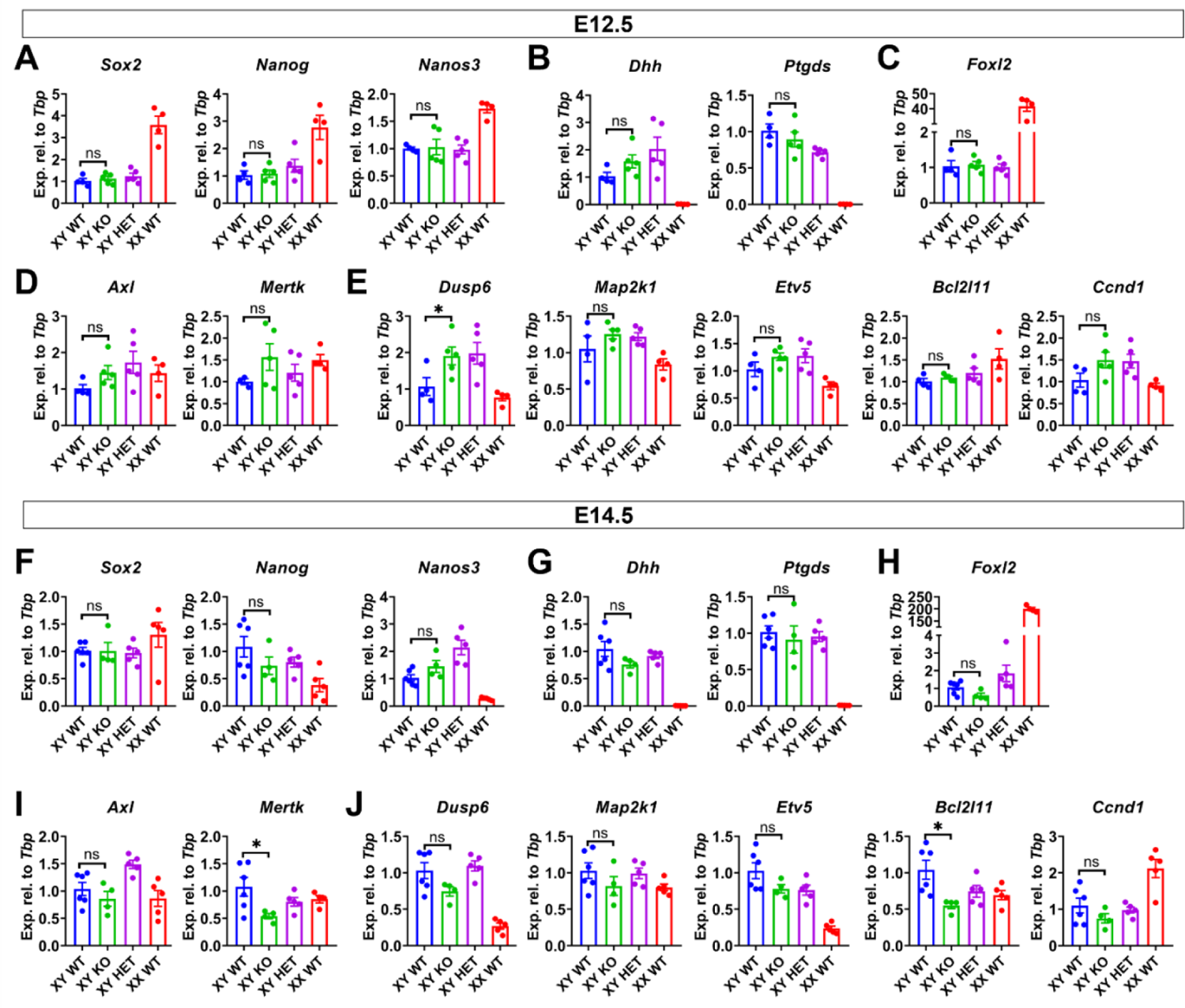
Gene expressions in E12.5 and E14.5 *Tyro3* knockout testes. At E12.5, qRT-PCR analysis of germ cell markers (**A**), SOX9 targets (**B**), an ovary marker (**C**), *Axl* and *Mertk* (**D**), and putative TYRO3 targets (**E**) in XY wildtype (XY WT, blue, n = 4), XY *Tyro3* knockout (XY KO, green, n = 5), XY heterozygous (XY HET, purple, n = 5) and XX wildtype (XX WT, red, n = 4). At E14.5, qRT-PCR analysis of germ cell markers (**F**), SOX9 targets (**G**), an ovary marker (**H**), *Axl* and *Mertk* (**I**), and putative TYRO3 targets (**J**) in (XY WT, blue, n = 6), XY *Tyro3* knockout (XY KO, green, n = 4), XY heterozygous (XY HET, purple, n = 5) and XX wildtype (XX WT, red, n = 5). Gene expression levels are normalized to *Tbp* and presented as relative expressions. Data is represented as Mean ± SEM. Unpaired Student’s *t* test. \**P* < 0.05; ns, not significant.

**Supplementary Figure 5.**
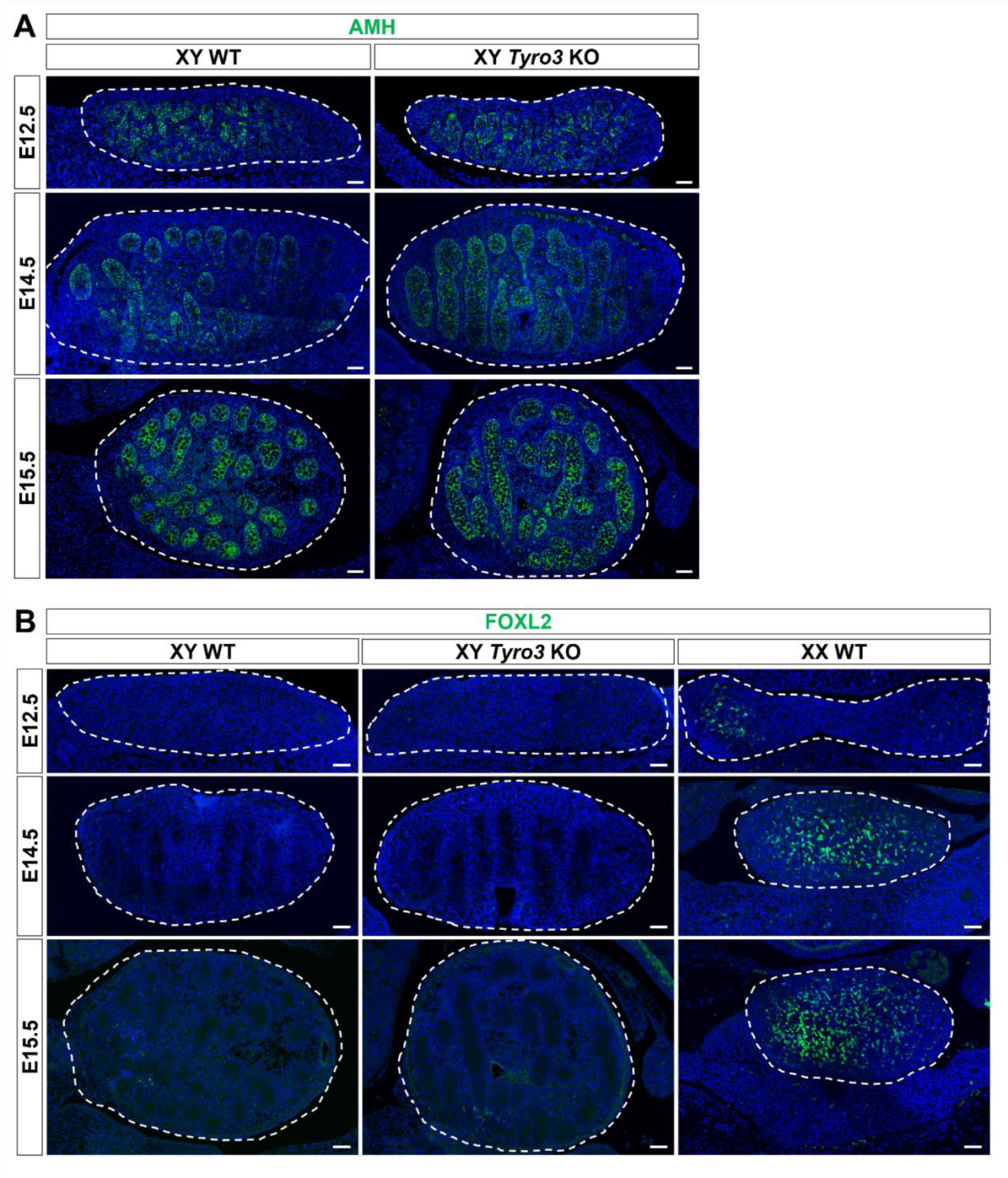
AMH and FOXL2 remain unchanged in *Tyro3* knockout testes. (**A**) Immunofluorescence staining for the Sertoli cell marker AMH (green) and (**B**) the female granulosa cell marker FOXL2 (green) at E12.5, E14.5 and E15.5. Nuclei are visualized with the nuclear marker DAPI (blue). Scale bar = 50 μm. Dashed lines outline gonads.

**Supplementary Figure 6.**
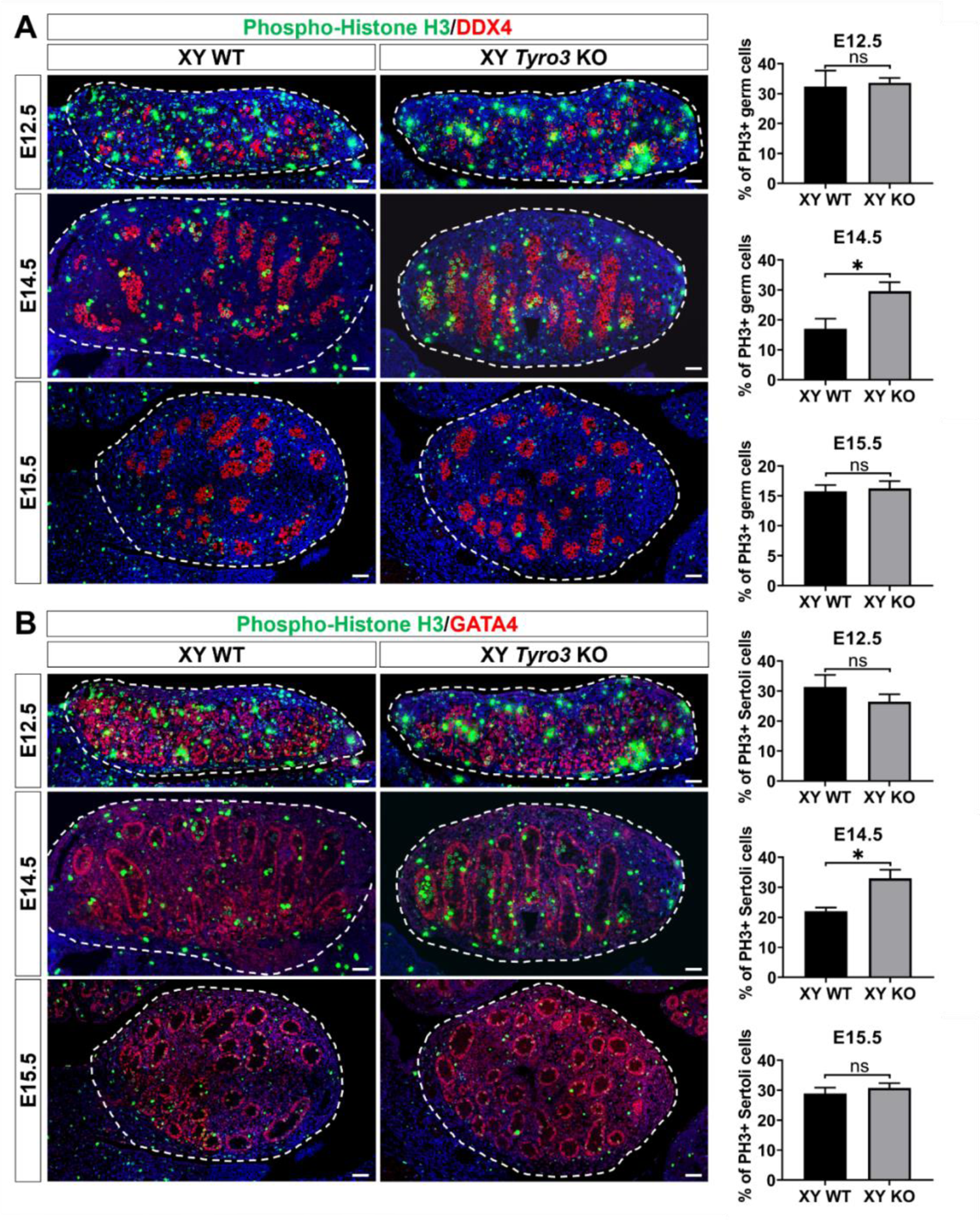
Cell proliferation in germ cells and Sertoli cells in *Tyro3* wildtype and knockout testes. (**A**) Immunofluorescence staining for the cell proliferation marker PH3 (green) and the germ cell maker DDX4 (red), and (**B**) the somatic maker GATA4 (red). Scale bar = 50 μm. Dashed lines outline gonads. Quantification is performed for PH3+ germ cells relative to the total germ cell population, and for PH3+ Sertoli cells relative to the total Sertoli cell population. At E12.5, n = 2 for XY WT, n = 3 for XY KO. At E14.5, n = 2 for XY WT, n = 3 for XY KO. At E15.5, n = 4 for XY WT, n = 5 for XY KO. Data is represented as Mean ± SEM. Unpaired two-tailed Student’s *t* test. \**P* < 0.05; ns, not significant.

**Supplementary Figure 7.**
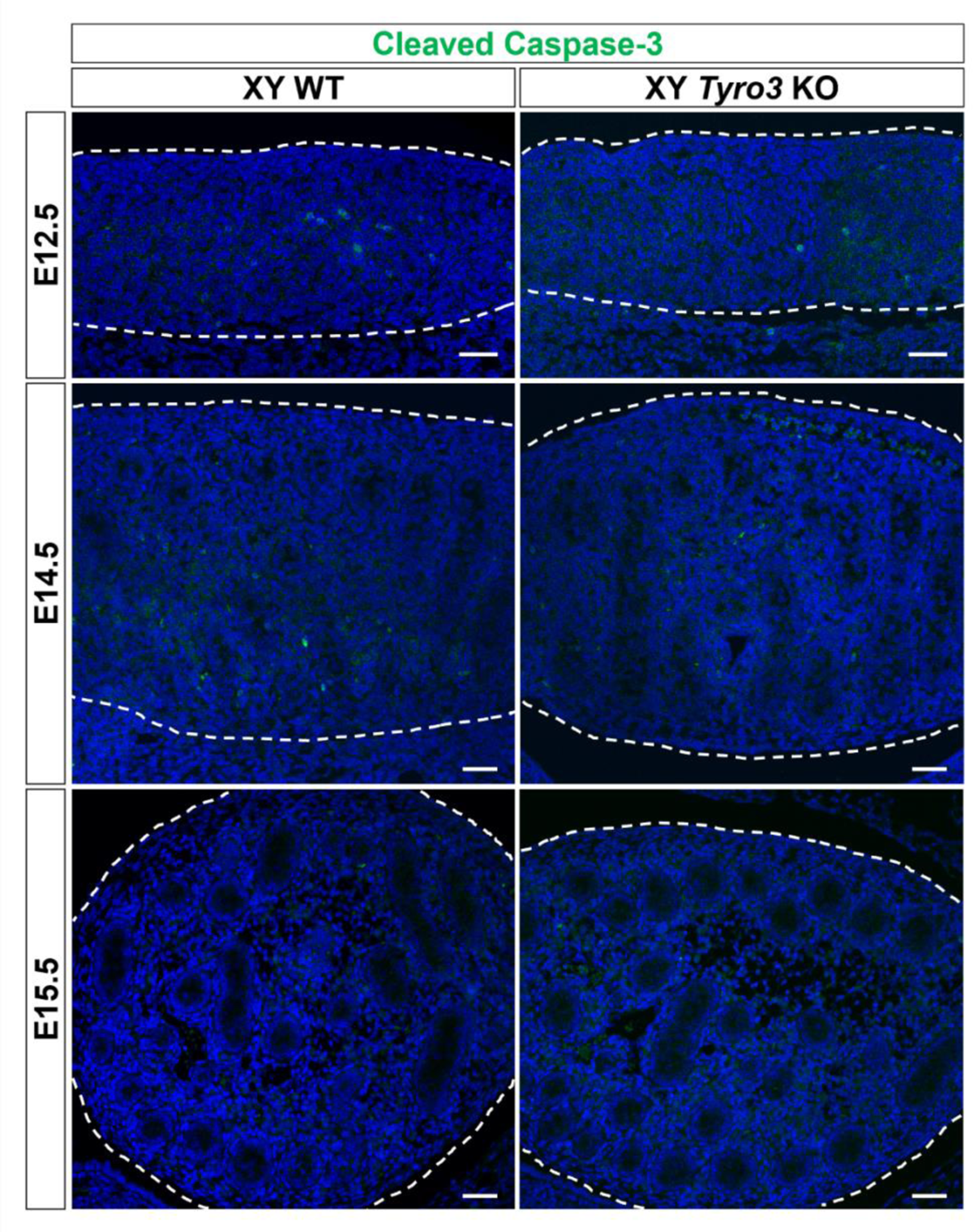
Absence of increased cell apoptosis in *Tyro3* knockout testes. Like in control gonads, immunofluorescence analysis targeting the cell apoptosis marker cleaved Caspase-3 (green) demonstrates very little cell apoptosis in *Tyro3* knockout testes at E12.5, E14.5 and E15.5. Nuclei are visualized with the nuclear marker DAPI (blue). Scale bar = 50 μm. Dashed lines outline gonads

## References

1. Koopman P, Gubbay J, Vivian N, Goodfellow P, Lovell-Badge R. Male development of chromosomally female mice transgenic for Sry. Nature 1991;351:117–21.

2. Sekido R, Bar I, Narváez V, Penny G, Lovell-Badge R. SOX9 is up-regulated by the transient expression of SRY specifically in Sertoli cell precursors. Developmental biology 2004;274:271–9.

3. Svingen T, Koopman P. Building the mammalian testis: origins, differentiation, and assembly of the component cell populations. Genes & development 2013;27:2409–26.

4. Chassot AA, Ranc F, Gregoire EP, Roepers-Gajadien HL, Taketo MM, Camerino G et al. Activation of beta-catenin signaling by Rspo1 controls differentiation of the mammalian ovary. Human molecular genetics 2008;17:1264–77.

5. Ottolenghi C, Pelosi E, Tran J, Colombino M, Douglass E, Nedorezov T et al. Loss of Wnt4 and Foxl2 leads to female-to-male sex reversal extending to germ cells. Human molecular genetics 2007;16:2795–804.

6. Bowles J, Knight D, Smith C, Wilhelm D, Richman J, Mamiya S et al. Retinoid signaling determines germ cell fate in mice. Science (New York, NY) 2006;312:596–600.

7. Durcova-Hills G, Capel B. Development of germ cells in the mouse. Current topics in developmental biology 2008;83:185–212.

8. Foster JW, Dominguez-Steglich MA, Guioli S, Kwok C, Weller PA, Stevanović M et al. Campomelic dysplasia and autosomal sex reversal caused by mutations in an SRY-related gene. Nature 1994;372:525–30.

9. Mansour S, Hall CM, Pembrey ME, Young ID. A clinical and genetic study of campomelic dysplasia. Journal of medical genetics 1995;32:415–20.

10. Wagner T, Wirth J, Meyer J, Zabel B, Held M, Zimmer J et al. Autosomal sex reversal and campomelic dysplasia are caused by mutations in and around the SRY-related gene SOX9. Cell 1994;79:1111–20.

11. Barrionuevo F, Bagheri-Fam S, Klattig J, Kist R, Taketo MM, Englert C et al. Homozygous inactivation of Sox9 causes complete XY sex reversal in mice. Biology of reproduction 2006;74:195–201.

12. Chaboissier MC, Kobayashi A, Vidal VI, Lützkendorf S, van de Kant HJ, Wegner M et al. Functional analysis of Sox8 and Sox9 during sex determination in the mouse. Development (Cambridge, England) 2004;131:1891–901.

13. Ming Z, Vining B, Bagheri-Fam S, Harley V. SOX9 in organogenesis: shared and unique transcriptional functions. Cellular and molecular life sciences: CMLS 2022;79:522.

14. Rahmoun M, Lavery R, Laurent-Chaballier S, Bellora N, Philip GK, Rossitto M et al. In mammalian foetal testes, SOX9 regulates expression of its target genes by binding to genomic regions with conserved signatures. Nucleic acids research 2017;45:7191–211.

15. Barrionuevo F, Georg I, Scherthan H, Lécureuil C, Guillou F, Wegner M et al. Testis cord differentiation after the sex determination stage is independent of Sox9 but fails in the combined absence of Sox9 and Sox8. Developmental biology 2009;327:301–12.

16. Jameson SA, Natarajan A, Cool J, DeFalco T, Maatouk DM, Mork L et al. Temporal transcriptional profiling of somatic and germ cells reveals biased lineage priming of sexual fate in the fetal mouse gonad. PLoS genetics 2012;8:e1002575.

17. Graham DK, DeRyckere D, Davies KD, Earp HS. The TAM family: phosphatidylserine sensing receptor tyrosine kinases gone awry in cancer. Nature reviews Cancer 2014;14:769–85.

18. Lemke G, Rothlin CV. Immunobiology of the TAM receptors. Nature reviews Immunology 2008;8:327–36.

19. Smart SK, Vasileiadi E, Wang X, DeRyckere D, Graham DK. The Emerging Role of TYRO3 as a Therapeutic Target in Cancer. Cancers 2018;10.

20. Lu Q, Gore M, Zhang Q, Camenisch T, Boast S, Casagranda F et al. Tyro-3 family receptors are essential regulators of mammalian spermatogenesis. Nature 1999;398:723–8.

21. Lu Q, Lemke G. Homeostatic regulation of the immune system by receptor tyrosine kinases of the Tyro 3 family. Science (New York, NY) 2001;293:306–11.

22. Xiong W, Chen Y, Wang H, Wang H, Wu H, Lu Q et al. Gas6 and the Tyro 3 receptor tyrosine kinase subfamily regulate the phagocytic function of Sertoli cells. Reproduction (Cambridge, England) 2008;135:77–87.

23. Matsubara N, Takahashi Y, Nishina Y, Mukouyama Y, Yanagisawa M, Watanabe T et al. A receptor tyrosine kinase, Sky, and its ligand Gas 6 are expressed in gonads and support primordial germ cell growth or survival in culture. Developmental biology 1996;180:499–510.

24. Blades F, Chambers JD, Aumann TD, Nguyen CTO, Wong VHY, Aprico A et al. White matter tract conductivity is resistant to wide variations in paranodal structure and myelin thickness accompanying the loss of Tyro3: an experimental and simulated analysis. Brain structure & function 2022;227:2035–48.

25. Jiménez A, Fernández R, Madrid-Bury N, Moreira PN, Borque C, Pintado B et al. Experimental demonstration that pre- and post-conceptional mechanisms influence sex ratio in mouse embryos. Molecular reproduction and development 2003;66:162–5.

26. Kasikara C, Davra V, Calianese D, Geng K, Spires TE, Quigley M et al. Pan-TAM Tyrosine Kinase Inhibitor BMS-777607 Enhances Anti-PD-1 mAb Efficacy in a Murine Model of Triple-Negative Breast Cancer. Cancer research 2019;79:2669–83.

27. Nurhayati RW, Ojima Y, Taya M. BMS-777607 promotes megakaryocytic differentiation and induces polyploidization in the CHRF-288-11 cells. Human cell 2015;28:65–72.

28. Paolino M, Choidas A, Wallner S, Pranjic B, Uribesalgo I, Loeser S et al. The E3 ligase Cbl-b and TAM receptors regulate cancer metastasis via natural killer cells. Nature 2014;507:508–12.

29. Bagheri-Fam S, Bird AD, Zhao L, Ryan JM, Yong M, Wilhelm D et al. Testis Determination Requires a Specific FGFR2 Isoform to Repress FOXL2. Endocrinology 2017;158:3832–43.

30. Al Kafri N, Hafizi S. Identification of signalling pathways activated by Tyro3 that promote cell survival, proliferation and invasiveness in human cancer cells. Biochemistry and biophysics reports 2021;28:101111.

31. Bagheri-Fam S, Ono M, Li L, Zhao L, Ryan J, Lai R et al. FGFR2 mutation in 46,XY sex reversal with craniosynostosis. Human molecular genetics 2015;24:6699–710.

32. Chen Y, Wang H, Qi N, Wu H, Xiong W, Ma J et al. Functions of TAM RTKs in regulating spermatogenesis and male fertility in mice. Reproduction (Cambridge, England) 2009;138:655–66.

33. DiNapoli L, Batchvarov J, Capel B. FGF9 promotes survival of germ cells in the fetal testis. Development (Cambridge, England) 2006;133:1519–27.

34. Bird AD, Frost ER, Bagheri-Fam S, Croft BM, Ryan JM, Zhao L et al. Somatic FGFR2 is Required for Germ Cell Maintenance in the Mouse Ovary. Endocrinology 2023;164.

35. Seervai RN, Wessel GM. Lessons for inductive germline determination. Molecular reproduction and development 2013;80:590–609.

36. Zhu S, Wurdak H, Wang Y, Galkin A, Tao H, Li J et al. A genomic screen identifies TYRO3 as a MITF regulator in melanoma. Proceedings of the National Academy of Sciences of the United States of America 2009;106:17025–30.

37. Li C, Scott DA, Hatch E, Tian X, Mansour SL. Dusp6 (Mkp3) is a negative feedback regulator of FGF-stimulated ERK signaling during mouse development. Development (Cambridge, England) 2007;134:167–76.

38. Prasad D, Rothlin CV, Burrola P, Burstyn-Cohen T, Lu Q, Garcia de Frutos P et al. TAM receptor function in the retinal pigment epithelium. Molecular and cellular neurosciences 2006;33:96–108.

39. Vollrath D, Yasumura D, Benchorin G, Matthes MT, Feng W, Nguyen NM et al. Tyro3 Modulates Mertk-Associated Retinal Degeneration. PLoS genetics 2015;11:e1005723.

40. Akalu YT, Mercau ME, Ansems M, Hughes LD, Nevin J, Alberto EJ et al. Tissue-specific modifier alleles determine Mertk loss-of-function traits. eLife 2022;11.

41. Bagheri-Fam S, Sim H, Bernard P, Jayakody I, Taketo MM, Scherer G et al. Loss of Fgfr2 leads to partial XY sex reversal. Developmental biology 2008;314:71–83.

